# Neural organization of speech production: A lesion-based study of error patterns in connected speech

**DOI:** 10.1101/544841

**Authors:** Brielle C Stark, Alexandra Basilakos, Gregory Hickok, Chris Rorden, Leonardo Bonilha, Julius Fridriksson

## Abstract

While numerous studies have explored single-word naming, few have evaluated the behavioral and neural correlates of more naturalistic language, like connected speech, which we produce every day. Here, in a retrospective analysis of 120 participants at least six months following left hemisphere stroke, we evaluated the distribution of word errors (paraphasias) and associated brain damage during connected speech (picture description) and object naming. While paraphasias in connected speech and naming shared underlying neural substrates, analysis of the distribution of paraphasias suggested that lexical-semantic load is likely reduced during connected speech. Using voxelwise lesion-symptom mapping (VLSM), we demonstrated that verbal (real word: semantically related and unrelated) and sound (phonemic and neologistic) paraphasias during both connected speech and naming loaded onto the left hemisphere ventral and dorsal streams of language, respectively. Furthermore, for the first time using both connected speech and naming data, we localized semantically related paraphasias to more anterior left hemisphere temporal cortex and unrelated paraphasias to more posterior left temporal and temporoparietal cortex. The connected speech results, in particular, highlight a gradient of specificity as one translates visual recognition from left temporo-occipital cortex to posterior and subsequently anterior temporal cortex. The robustness of VLSM results for sound paraphasias derived during connected speech was notable, in that analyses performed on sound paraphasias from the connected speech task, and not the naming task, demonstrated significant results following removal of lesion volume variance and related apraxia of speech variance. Therefore, connected speech may be a particularly sensitive task on which to evaluate further lexical-phonological processing in the brain. The results presented here demonstrate the related, though different, distribution of paraphasias during connected speech, confirm that paraphasias arising in connected speech and single-word naming likely share neural origins, and endorse the need for continued evaluation of the neural substrates of connected speech processes.

## Introduction

Much of what we know about the neural architecture of language production is grounded in studies of single word retrieval (Levelt, 1999; Foygel and Dell, 2000; Walker and Hickok, 2016). To successfully retrieve a word, such as the name of an object, one must access at least three levels of information: conceptual (what it is), lexical (its associated word) and phonological (the sounds to select and organize). When we describe a situation, however, we do not simply name objects. Instead, we select two or three words per second from an active vocabulary spanning an estimated 40,000 words (Levelt, 1989), and making this feat more complex, our word selection is vulnerable to “competition” from multiple sources. For example, errors in connected speech tend to be phonologically or semantically related word substitutions or omissions, the replacement of the intended target word with a word that we intended to use in a future production, or the perseveration on a word that was used recently (Lashley, 1951; Garrett, 1975). Selecting a word in connected speech, therefore, requires a delicate balance of target word activation and suppression of alternatives, where alternatives come in the form of previously selected forms, subsequent forms in various stages of planning, and similar competitor forms, all coordinated across multiple hierarchical levels. Therefore, connected speech stands in contrast to single word production, where one’s full attention is focused on the production of a single word rather than the selection and inhibition of many lexical items.

While models explaining single word retrieval have been developed and studied heavily, relatively few models have focused on modeling the complex lexical retrieval system engaged during sentence production (Garrett, 1975; Dell, 1986; Chang, 2002; Chang *et al.*, 2006). Indeed, connected speech production likely depends on cognitive operations that extend beyond lexical processing (Fergadiotis and Wright, 2015) alongside a lexical retrieval system where activation of word forms is dynamically changing across the course of the utterance. One step in evaluating the lexical retrieval processes during connected speech has been through the use of blocked cyclic picture naming, where subjects repeatedly name pictures in sets of either semantically related or unrelated pictures, cycling several times through the pictures in each block. While blocked cyclic picture naming is not true connected speech, it provides a multiword environment more complex than naming. These studies have found a semantic interference effect, such that naming semantically related pictures becomes increasingly slower when repeatedly naming from a semantic category even when several unrelated trials occur (Dell *et al.*, 2000; Oppenheim *et al.*, 2010; Chang *et al.*, 2012; Goldrick and Larson, 2016; Hughes and Schnur, 2017). Semantic interference may result from lexical competition, such that competition grows more intense when retrieving words in a homogeneous context because of the growing number of competitors (i.e. semantically related words) but semantic interference may likewise result from impaired inhibition, such that lexical competitors cannot be suppressed (Oppenheim *et al.*, 2010). It is an open question whether work on paradigms such as blocked cyclic picture naming will generalize to more natural speech.

A handful of prior studies have evaluated neurogenic communication disorders, such as post-stroke aphasia, to compare the distribution and properties of word-level errors (paraphasias) during connected speech and single word retrieval. It seems intuitive that the underlying processes involved in producing a paraphasia on the two tasks should be similar, and for the distribution of such errors on one task to predict the distribution on the other. However, there are reports of patients with aphasia demonstrating superior word retrieval during confrontation naming compared to connected speech (Manning and Warrington, 1996; Schwartz and Hodgson, 2002; Wilshire and McCarthy, 2002) and vice versa (Zingeser and Berndt, 1990; Ingles and Mate-Kole, 1996; Pashek and Tompkins, 2002). It is therefore becoming increasingly evident that connected speech requires dynamic changes in the linguistic system, resulting in differing distributions of linguistic components compared to naming. An early study evaluated word retrieval deficits in picture naming (the Boston Naming Test) and word-finding difficulty for nouns during connected speech production and found a ∼58% overlap in variance between the two variables (Nicholas *et al.*, 1989). A later study evaluated naming performance and connected speech in 14 participants with mild and moderate aphasia, finding that there was superior word retrieval and overall more self-corrected errors during the connected speech task and that naming ability was not significantly correlated with connected speech paraphasias (Mayer and Murray, 2003). Further evaluation by other groups suggested that naming ability may relate more to the ability to retrieve nouns, in particular, during everyday conversation (Herbert *et al.*, 2008). Further, a recent study in 98 participants with aphasia found that picture naming ability was not predictive of the proportion of paraphasias made during connected speech, finding that paraphasias produced in discourse in isolation were not well predicted by performance on confrontation naming tests (Fergadiotis and Wright, 2015). Taken together, these studies encourage the direct comparison of the distribution of paraphasias between connected speech and naming for improved understanding of the dynamic lexical retrieval system employed during natural speech. Explanations regarding the reason for the discrepancy of paraphasia distributions between connected speech and single word retrieval have revolved around the differential context-dependent, non-linguistic demands required by connected speech as well as by linguistic factors such as the effects of lexical neighborhood and phonemic similarity (Foygel and Dell, 2000; Penn, 2000; Pashek and Tompkins, 2002).

If indeed there is disparity between the distributions of paraphasias in connected speech and naming, it raises the question regarding the extent to which neural mechanisms underlying single-word paraphasias during connected speech and confrontation naming are similar. Few studies have evaluated the neural correlates of paraphasias made during connected speech. In 70 participants with primary progressive aphasia, Wilson and colleagues associated phonemic paraphasias made during a connected speech task with atrophy of the left superior temporal gyrus (Wilson *et al.*, 2010). Meanwhile, studies of naming in post-stroke aphasia have localized phonemic paraphasias and speech production errors to dorsal stream structures (Schwartz *et al.*, 2012; Dell *et al.*, 2013). Whilst the methodology used, and the population studied across these examples, are not directly comparable, the results of these investigations encourage further evaluation of the neural structures underlying connected speech compared with naming paraphasias.

In the current study, we evaluated the distribution of paraphasias and associated brain damage during a connected speech task and a naming task. This work contributes to the extension of single word and sentence models to natural speech.

## Materials and Methods

### Participants

120 individuals with chronic (>6 months) single-event left hemisphere stroke were retrospectively included in this study. These participants were recruited as part of larger studies conducted to evaluate treated aphasia recovery and lesion-deficit relationships. Participants were recruited between 2007 and 2017 at the University of South Carolina or Medical University of South Carolina, where all studies had IRB approval. Inclusion parameters for our studies include a single left hemisphere stroke, at least six months post-stroke and an age limit of 18 – 85 years. Exclusion parameters for our studies include bilateral stroke; right hemisphere stroke; brainstem stroke; non-proficiency in English and pre-morbid left handedness. All participants gave written consent for study inclusion. Language evaluations were conducted within a week of neuroimaging evaluations. As this was a retrospective study, no part of the study procedures or analyses was preregistered prior to the research being undertaken.

Participants were from different studies done within our lab and therefore completed a variety of language assessments. In this case, participants had either a naming assessment, a connected speech assessment or both assessments. In the following section, we will describe these participants as belonging to the CS task group (connected speech) or the PNT task group (naming).

We originally identified 114 participants with a naming assessment and 63 with a connected speech assessment; 36 participants had both assessments. To create two mutually exclusive groups with which to compare lesion data, we matched participants with a naming assessment to those with a connected speech assessment using case-control matching in SPSS 25. We matched on aphasia severity (from the Western Aphasia Battery [WAB] Aphasia Quotient [AQ] from Kertesz, 2007) using a FUZZY interval of one standard deviation of the CS task group’s AQ. This resulted in two mutually exclusive groups: a connected speech task group of N=63 (CS task group) and a naming task group of N=57 (PNT task group).

#### Description of Task Groups

##### Connected Speech (CS) Task Group (N=63)

This task group comprised participants who had completed a connected speech assessment, which consisted of describing a set of three pictures (described in *Behavioral Tasks*). There were 63 participants (40 males; age, M=59.11±10.05 yrs; months post-stroke, M=40.11±45.9) included in this subset with the following aphasia types: anomic, 12; Broca’s, 19; conduction, 8; global, 1; transcortical sensory, 1; and Wernicke’s, 2. Twenty participants scored above the cut-off for demonstrating aphasia (>93.8 out of 100), as specified by the WAB AQ. Average AQ was 75.12±23.82 and lesion volume was 92.99±84.81 cm^3^.

##### Naming (PNT) Task Group (N=57)

This task group included those participants who completed a visual confrontation naming task involving the naming of objects, the Philadelphia Naming Test (Roach *et al.*, 1996), described below in *Behavior Tasks*. There were 57 participants (32 males; age, M=61.68±12.02 yrs; months post-stroke, M=31.91±37.16) included in this subset with the following aphasia types: anomic, 25; Broca’s, 19; conduction, 6; global, 1 and Wernicke’s, 2. Four participants scored above the cut-off for demonstrating aphasia (>93.8 out of 100), as specified by the WAB AQ. Average AQ was 68.78±23.91 and lesion volume was 111.27±94.44 cm^3^.

These groups are described and statistically compared in Table 1A.

**Table 1.**
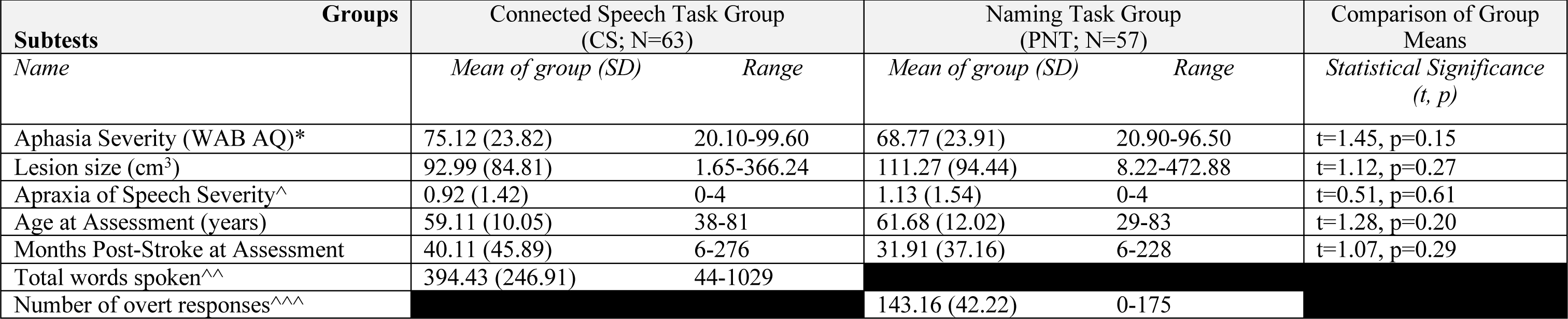
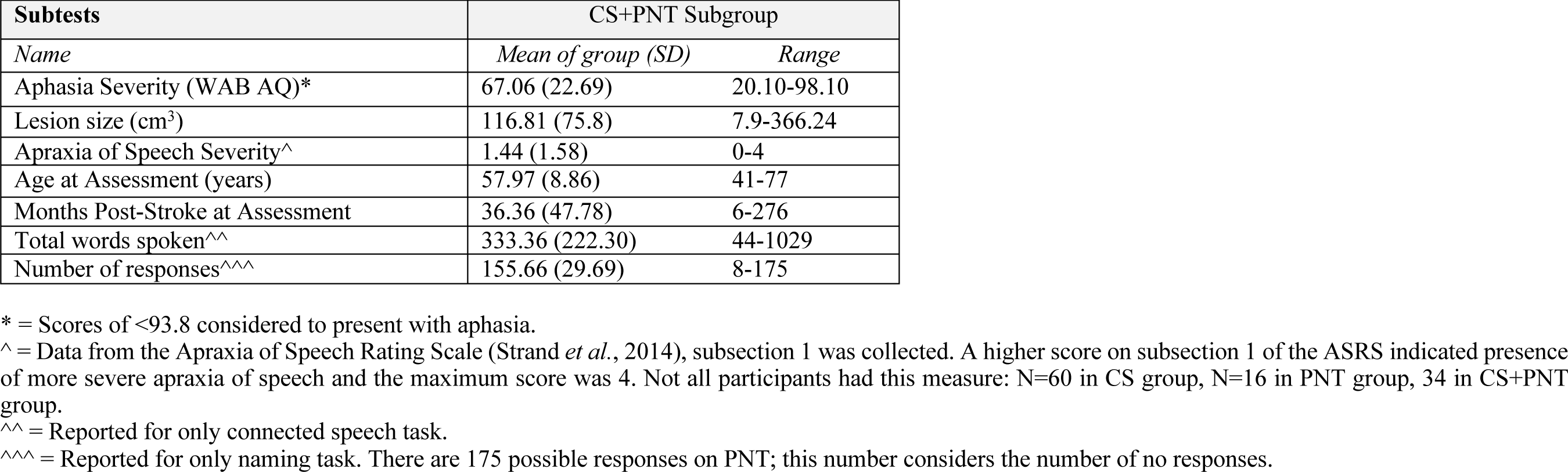
<u>Demographics of all participants (N=120) included in the study</u> Table 1A: Demographics of the mutually exclusive task groups on which the VLSM analysis was completed. A comparison using an independent samples t-test (p<0.05) is shown in the right-right column, comparing the CS and PNT task groups. Table 1B: Demographics of the CS+PNT subgroup participants (N=36) on which the behavioral analysis was completed.

We used these two mutually exclusive naming and connected speech groups to compare lesion data. A subset of participants had *both* a naming and connected speech assessment; we used this group to directly compare the distribution of paraphasias across tasks. This group is described, below.

##### Connected Speech and Naming (CS+PNT) Subgroup

A subgroup of participants who completed *both* the naming and connected speech tasks were identified (N=36, 25 males). All members of this group were members of the CS task group used in the lesion data analysis; they were not members of the PNT task group. Aphasia types in this group were: anomic, 9; Broca’s, 17; conduction, 5; and Wernicke’s, 2. Three participants scored above the WAB AQ cut off of 93.8. The average WAB AQ was 67.06±22.69 and lesion volume was 116.81±75.8 cm^3^. Table 1B presents demographics from this subgroup.

A flowchart of the groupings described above is available in Supplementary Figure 1.

**Figure 1:**
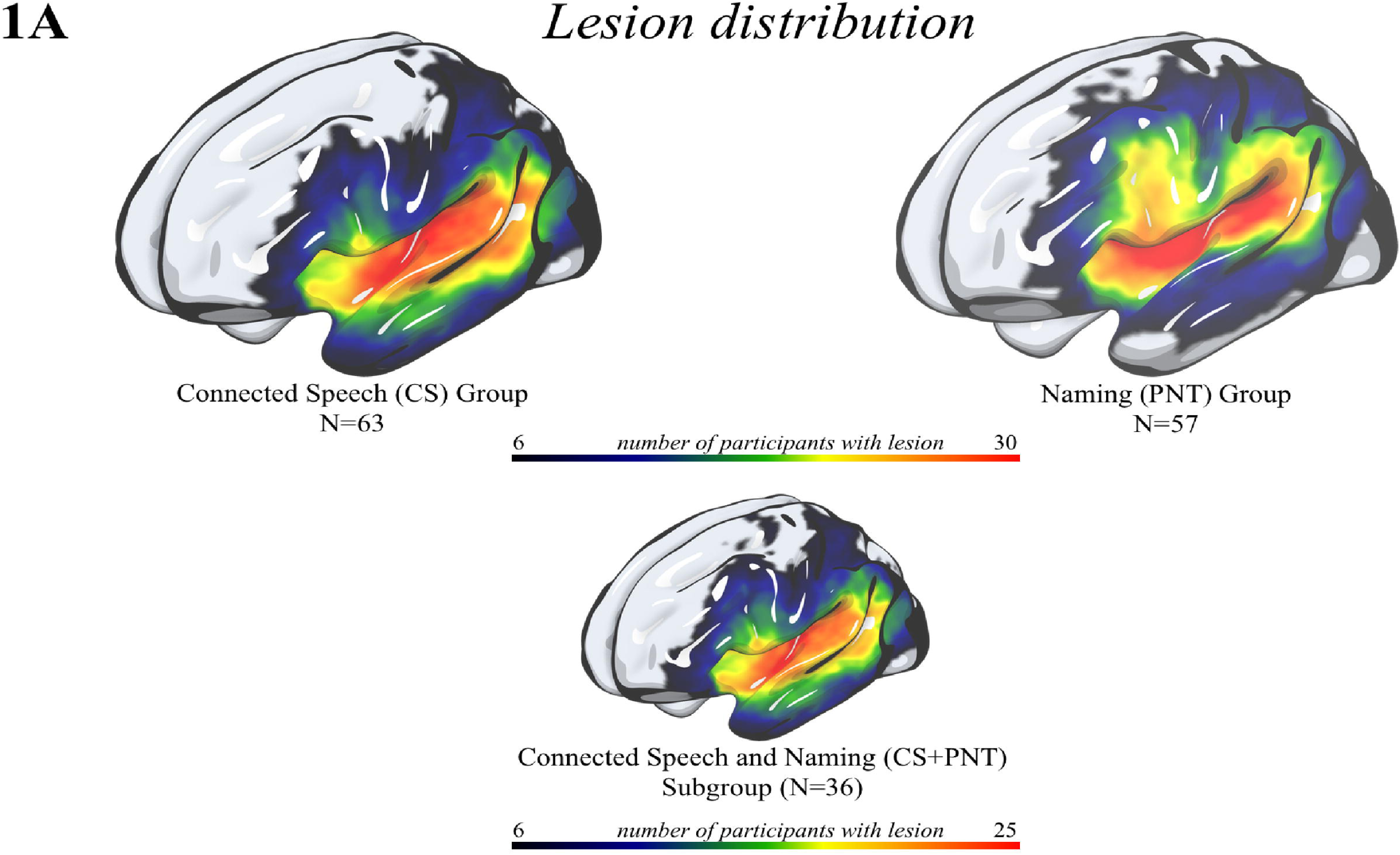
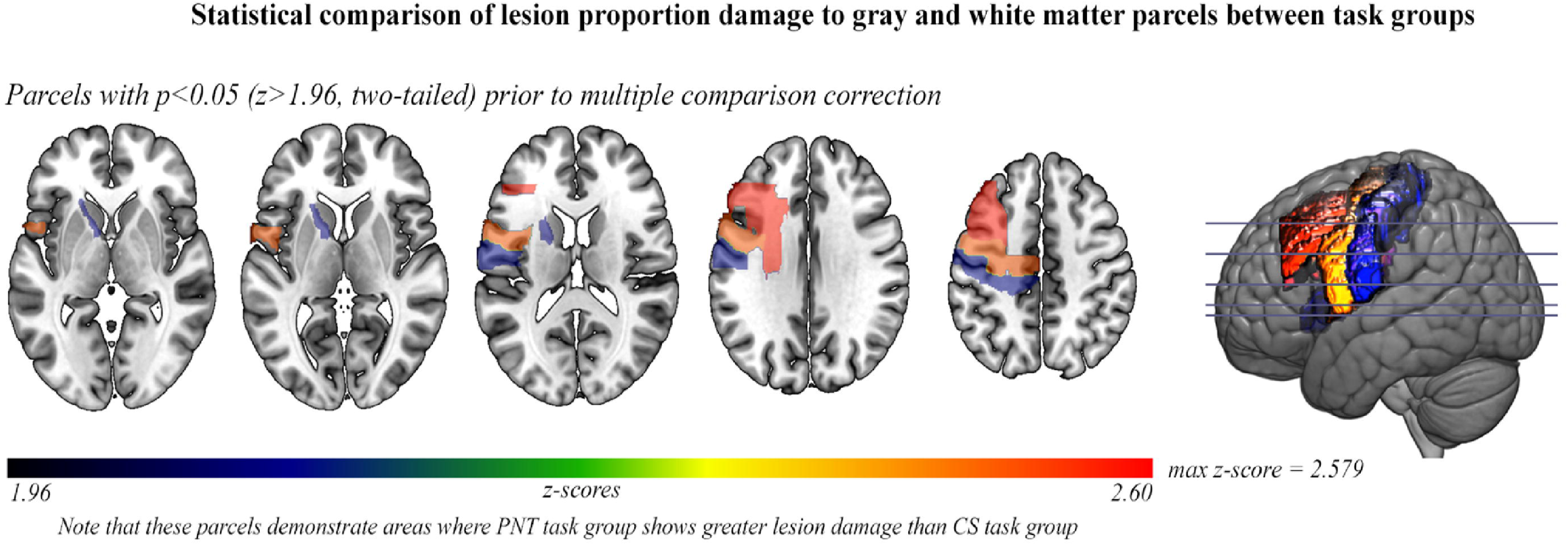
<u>Distribution of lesions for each task group</u> (CS task group, PNT task group, CS+PNT subgroup). 1A: A heat map on rendered surface showing lesion distributions for the participants included in the CS task group (top, left); PNT task group (top, right) and CS+PNT subgroup (bottom). Areas in red demonstrate where more participants share a lesion to that voxel. 1B: Prior to multiple comparison correction, a comparison of left hemisphere lesion proportion damage to parcels between task groups shown on axial slices with a left hemisphere sagittal cross-slice. Parcellation was from the Johns Hopkins University atlas (Faria et al., 2012), which contained gray- and white-matter parcels (182 in the whole brain). When comparing lesion damage to each parcel between the CS and PNT task groups, there was not a significant difference between lesion proportion damage to a parcel after multiple comparison correction (FDR; q=0.05). Prior to multiple comparison correction, six dorsal stream parcels were significantly more damaged in the PNT task group than the CS task group at p<0.05 (z>1.96); these are shown in the figure. P-values for these parcels ranged from p=0.0098 (middle frontal gyrus) to p=0.036 (underlying white matter, internal capsule.

### Behavioral Tasks

#### Connected Speech Task: Picture Description

Participants described three pictures from speech/language test batteries (Dabul, 2000; Goodglass *et al.*, 2000; Kertesz, 2007) for ∼2 minutes each while being audio-video recorded. Participants were instructed to describe the pictures using complete sentences whenever possible.

##### Motivation

We used picture description as a measure of connected speech because picture description provides clear conceptual targets which make identifying the type of paraphasia, and its intended target, more reliable. For instance, in describing a picture of a circus, we can make the reliable judgment that the word ‘cat’ is a semantically related paraphasia of ‘lion.’ Furthermore, utterance context in picture description provides a source of constraint that is not available in picture naming. Specifically, because connected speech provides context (which naming does not), which may elicit the production of verbs, adjectives, adverbs, and other referential words and phrases, we are more often than not able to identify the targets of paraphasias. For example, while describing a circus scene, one participant said: “the clown is sitting on its back legs,” erroneously saying ‘clown’ instead of ‘lion.’ Because we knew the context of the scene, ‘clown’ and ‘lion’ were easily classified as semantically related (more specifically, thematically semantically related).

##### Transcription

Speech was transcribed from audiovisual recordings by a speech-language pathology Master’s student who also had a master’s degree in linguistics (KK). Using CHAT/CLAN software, a transcription and analysis software for speech (MacWhinney, 2000), each transcribed utterance was time-stamped to its appropriate place on the audiovisual media. After initial transcription, author BS then simultaneously watched and read each transcript; discrepancies were discussed and corrected. Both author BS and transcriber KK were blinded to participant AQ and lesion site/volume at the time of transcription and transcription checking. Transcriptions from all three picture description attempts were combined for each participant; two participants described only one picture and seven described two pictures.

##### Coding Paraphasias

Transcriptions were coded using CHAT/CLAN program parameters (MacWhinney, 2000). CHAT is a software program which allows for tags to be added to words, phrases and utterances, whilst CLAN is its component analysis program. Paraphasias were coded by hand using CHAT format, such that individual words were tagged as a verbal paraphasia (two types: semantically related and unrelated), a phonemic paraphasia or a neologism (see operational definitions, below). CLAN was then used to extract the number and type of paraphasias in each transcript based on these tags.

Verbal paraphasias were coded as follows: a ‘semantically related’ paraphasia was a real word that was related to a known target (‘cat’ for ‘dog’) and an ‘unrelated’ paraphasia was a real word that was otherwise unrelated to a target (e.g. ‘joke’ for ‘boy’) or a real word of an unknown relation to the target. Paraphasias were coded as phonemic using pre-determined parameters for the number of shared phonemes of the production with the target. Briefly, phonemic paraphasias refer to the substitution of a word with a nonword that preserves at least half of the segments and/or number of syllables of the intended word. The specific criteria for identifying a phonemic paraphasia were as follows. For one-syllable words, consisting of an onset (initial phoneme or phonemes) plus vowel nucleus plus coda (final phoneme or phonemes), the error must match on two out of three elements: onset plus vowel nucleus, vowel nucleus plus coda or onset plus coda. The part of the syllable that is in error may be a substitution, addition, or omission. For one-syllable words with no onset (e.g., eat) or no coda (e.g., pay), the absence of the onset or coda in the error was also counted as a match. For multi-syllabic words, the error must have complete syllable matches on all but one syllable, and the syllable with the error must meet the one-syllable word match criteria stated above. Neologisms were identified as non-words which did not fit phonemic paraphasia criteria. Other groups have further subdivided neologisms into abstruse and phonemically-related neologisms (Walker *et al.*, 2018), but we did not do so here.

##### Interrater reliability of error coding

Authors BS and AB, both blinded to lesion site/volume, aphasia types and WAB AQ scores, performed reliability for error coding on 20% of the transcripts. Paraphasia coding agreement was subsequently compared between BS, AB and the original coder (KK). Compared with KK’s paraphasia coding, AB showed 94% agreement and BS showed 95% agreement. AB and BS agreed on 98.63% of samples.

#### Naming Task: Philadelphia Naming Test (PNT)

The PNT required participants to verbally name 175 black-and-white pictures; answers ranged in length from one to four syllables (Roach *et al.*, 1996). If more than one response was given for a single picture, the last response was scored. Paraphasias (semantically related; unrelated; phonemic; neologism) were identified in the same way as they were identified in the connected speech task (see above). Correct responses, self-corrections, mixed errors, perseverations and articulation errors were also recorded but were not used here. ‘No response’ trials, where a participant did not attempt to name the picture, were documented.

### Analysis of Behavioral Data

#### Distribution of Paraphasias

We calculated the proportion of paraphasias (i.e. the proportion of semantically related paraphasias per all attempts) for both connected speech and naming tasks. We calculated the proportion of each type of paraphasia on the PNT as: (# paraphasias / (175-no responses)). As the PNT comprises only objects (nouns), we wanted to provide a comparable calculation for the connected speech task. We determined that the majority of paraphasias replaced noun targets (Supplementary Table 1), and we therefore calculated the proportion of each type of paraphasia on the connected speech tasks as: (# paraphasias / total number of nouns).

##### Identifying the parts of speech of paraphasia targets during connected speech

Author BS classified all paraphasia targets as a noun, verb, another part of speech (adjective, adverb, etc.), or as unknown. On the same sample used for interrater reliability of error coding (above; 20% of sample), authors AB and GH separately classified the intended target’s part of speech. We subsequently compared this classification across four categories (noun [including pronoun]; verb; other [adjective, adverb, etc.] and unknown). All raters were blinded to lesion volume and site as well as AQ. Across the categories of parts of speech, the agreement between BS, AB and GH was 90.37% (nouns, 88.19%; verbs, 95.83%; other, 90.97% and unknown, 86.46%). As there was good agreement in coding part of speech, and overall agreement that targets being replaced by paraphasias were largely nouns, we opted to calculate the proportion of paraphasias as a proportion of total nouns. Supplementary Table 1 shows the classification of parts of speech for each paraphasia type by author BS.

##### Analysis

We only compared the distribution of paraphasias in the CS+PNT subgroup, because members of this group had completed both a connected speech and a naming assessment. In this group, we performed multinomial logistic regression with type of task as the dependent variable (naming, connected speech) with the proportion of paraphasias of each subtype (phonemic, neologistic, semantically related, unrelated) as the covariates.

### Neuroimaging Data Acquisition and Processing

#### Image Acquisition

The 3T scanner was upgraded during data collection. Earlier structural images were collected using a 12-channel head coil on a Siemens Trio scanner while later images were collected using a 20-channel head coil on the Prisma Fit scanner. The Trio scans were acquired with the following parameters. T1-weighted image was acquired utilizing MP-RAGE sequence with 1 mm isotropic voxels, a 256 x 256 matrix size, and a 9-degree flip angle. We used a 192-slice sequence with TR=2250 ms, TI=925 ms and TE=4.11 ms. T2-weighted image was acquired using a sampling perfection with application optimized contrasts using a different flip angle evolution (3DSPACE) sequence. This 3D TSE scan used a TR = 3200 ms, a TE of 567 ms, variable flip angle and a 256 x 256 matrix scan with 160 slices (1 mm thick) using parallel imaging (GRAPPA=2, 80 reference lines). The T1-weighted images from the 3T Prisma scans were acquired using the acquisition parameters described above. T2-weighted images were acquired using 3DSPACE sequence, using identical parameters to the Trio sequence but with 176 slices (1 mm thick). For each individual, the same volume center and slice angulation parameters were used for the T1 and T2 sequences.

#### Image Processing

Lesions were manually drawn on the T2-weighted image by a neurologist (author LB), who was blinded to the participant’s language scores at the time of the lesion drawing. The T2 image was co-registered to the T1 image, and these parameters were used to re-slice the lesion into the native T1 space. The resliced lesion maps were smoothed with a 3mm full-width half maximum Gaussian kernel to remove jagged edges associated with manual drawing. We then performed enantiomorphic segmentation-normalization (Nachev *et al.*, 2008) using SPM12 and MATLAB scripts we developed (Rorden *et al.*, 2012) as follows: first, a mirrored image of the T1 image (reflected around the midline) was created, and this mirrored image was co-registered to the native T1 image. We then created a chimeric image based on the native T1 image with the lesioned tissue replaced by tissue from the mirrored scan (using the smoothed lesion map to modulate this blending). SPM12’s unified segmentation-normalization (Ashburner and Friston, 2005) was used to warp this chimeric image to standard (MNI) space, with the resulting spatial transform applied to the native T1 image as well as the lesion map and the T2 image (which used the T1 segmentation parameters to mask non-brain signal). The normalized lesion map was then binarized, using a 50% probability threshold. Figure 1A shows the overlap of lesions for our sample of participants, separated by task group.

### Neuroimaging Data Analysis

#### Lesion-symptom Mapping

To evaluate lesion damage significantly associated with word-finding errors in connected speech and naming, we conducted univariate voxel-based lesion-symptom mapping (VLSM; Bates et al., 2003). This analysis was conducted on the two task groups: CS and PNT. We used the proportion of each type of paraphasia (neologistic, phonemic, semantically related, unrelated) as the dependent variable. Voxelwise data were analyzed at p<.05 (one-tailed) with 5000 permutations (Winkler et al., 2014). Cluster thresholding was then employed (z>2.33, p<0.01) and corrected for multiple comparisons using nonparametric permutation (5000 permutations; Winkler et al., 2014). Specifically, the order of operations for this analysis was as follows: 1) compute voxelwise statistics for 5000 permutations, 2) zero all voxels less than z=1.645 (p>0.05), 3) measure the number of voxels in the largest surviving cluster and save this variable, 4) rank order the 5000 observations of max cluster size, 5) threshold at the 100^th^ largest cluster size (i.e. given random data, we observed only a single cluster of this size in 1% of the permutations) and 6) zero all voxels less than z=1.645 and zero all clusters smaller than the cluster identified in step #5. This cluster thresholding is part of the VLSM software detailed in Wilson *et al.*, (2010) as well as in our own program, NiiStat (https://www.nitrc.org/projects/niistat/).

VLSM analyses were conducted on voxels in which at least 10% of participants shared a lesion. To correct for lesion volume, we used Freedman-Lane permutation correction with lesion volume as a nuisance variable (Winkler et al., 2014). As some paraphasias significantly correlated with apraxia of speech (AOS) severity in the connected speech condition, we also regessed AOS severity (a score of 0 to 4 where 0 means no AOS; scores derived from the Apraxia of Speech Rating Scale, ASRS Strand et al., 2014) from these error types using Freedman-Lane permutation.

#### Comparison of neural data between mutually exclusive groups

The goal of this study was to compare the results of a voxelwise lesion-symptom mapping (VLSM) analysis between the two task groups. A basic assumption regarding VLSM is that results will generalize beyond the specific participants included in the analysis. There are two possible challenges for this assumption when comparing two distinct groups of subjects: 1) that lesion distribution is significantly different between task groups, and 2) that the number of participants with lesion damage per parcel is significantly different between task groups. We addressed these assumptions, below, and provided a supplementary table (Supplementary Table 2) of lesion proportion damage and number of participants with damage to each parcel.

**Table 2:** <u>Distribution of paraphasia types by task</u>. The CS (connected speech) and PNT (naming) task groups are shown on the left. The CS+PNT subgroup, comprised of individuals who had both a connected speech and naming assessment, is shown on the right. Recall that the percentage of paraphasias was calculated a proportion of total nouns for the connected speech task and as a proportion of total overt attempts for the naming task. Significant difference between the distributions of paraphasias during connected speech and naming was calculated only for the CS+PNT subgroup, as this was a direct comparison between individuals who completed both tasks. *\* Indicates a significant difference from the multinomial logistic regression analysis performed on the CS+PNT subgroup.*

| *Task Groups* | **CS Task Group**<br>M (SD) % | **PNT Task Group**<br>M (SD) % | **CS+PNT subgroup**<br>M (SD) % |  |  |
| --- | --- | --- | --- | --- | --- |
| *Task Types* | Connected Speech | Naming | Connected Speech | Naming | Multinomial logistic regression significance |
| *Paraphasia Types* |  |  |  |  |  |
| Neologistic | 6.03 (11.42) | 6.89 (19.04) | 9.00 (13.75) | 6.97 (15.50) | p=.13 |
| Phonemic | 6.49 (8.93) | 4.69 (7.77) | 8.22 (9.66) | 9.06 (9.90) | p=.48 |
| Semantically Related | 0.72 (1.21) | 8.89 (9.94) | 0.83 (1.25) | 8.89 (8.28) | p=.001** |
| Unrelated | 3.27 (5.38) | 3.26 (5.10) | 4.75 (5.80) | 2.50 (3.63) | p=.015* |

1. Lesion distribution was not significantly different between task groups, which we demonstrated at both the voxel- and parcel-level (gray- and white-matter parcels derived from the Johns Hopkins University [JHU] atlas, [Faria et al., 2012]). There was no significant difference in proportion damage at the voxel-level between task groups (z-score range: −3.34 - 4.10), which was calculated using a two-tailed Liebermeister test (p<0.05) and FDR correction for multiple comparison (Rorden *et al.*, 2007) in all left hemisphere voxels where at least 10% of people had damage. Further, there was no significant difference in lesion damage to any parcel between task groups (z-score range: - 1.80-2.579), which was calculated using a two-tailed t-test (p<0.05) and FDR correction for multiple comparison in all left hemisphere parcels where at least 10% of people had damage. Prior to multiple comparison correction, six parcels had a score of z>1.96 (therefore, where the PNT task group had more lesion proportion damage than the CS task group): a posterior section of the middle frontal gyrus (two-tailed z=2.579, p=0.0099), white matter (superior corona radiata) (two-tailed z=2.572, p=0.01), precentral gyrus (two-tailed z=2.517, p=0.012), postcentral gyrus (two-tailed z=2.127, p=0.033) and white matter (anterior limb of internal capsule) (two-tailed z=2.099, p=0.036). Figure 1B displays these parcels. However, given no statistical significance after multiple comparison correction, we stipulate that lesion distribution was not significantly different between task groups.
2. The number of participants with lesion damage to a parcel was not significantly different between task groups. We analyzed the number of participants with damage to a parcel, compared across all left hemisphere parcels (gray- and white-matter, JHU atlas) in which at least 10% of subjects had damage. A two-tailed t-test (p<0.05) rejected the null hypothesis, t(137)=0.846 (p=0.399), providing evidence that there was not a significant difference in the number of participants with damage to a left hemisphere parcel between task groups.

This evidence suggests that the task groups are well-matched and differences between them accounted for and, as such, comparisons between groups must be acceptable under the assumption that VLSM results should generalize.

##### Posthoc analyses

Common mechanisms were tested using a conjunction analysis (p<0.05) on significant clusters identified by the VLSM (Nichols *et al.*, 2005), showing clusters of voxels that, when damaged, associated with paraphasias in both the PNT and the CS task groups (Nichols *et al.*, 2005). Therefore, by examining the supra-threshold voxels shared across task group analyses, we can determine whether damage to a brain region underlies a specific paraphasia type for just one task (e.g. connected speech) or both tasks (e.g. naming and connected speech).

## Results

### Behavioral

#### Paraphasia Profiles in CS and PNT Task Groups

Recall that for the CS task group, the proportion of paraphasias was calculated as a proportion of total nouns, whereas for the PNT task group, the proportion of paraphasias was calculated as a proportion of total attempts. The distribution of paraphasias for each group is shown in Table 2.

For each task group, and for the CS+PNT subgroup, we computed a correlation matrix (Figure 2). This matrix included paraphasias made during the respective task; biographical variables (age, time since stroke, lesion volume); and language variables (WAB AQ, total words spoken / total items named, apraxia of speech severity score).

**Figure 2:**
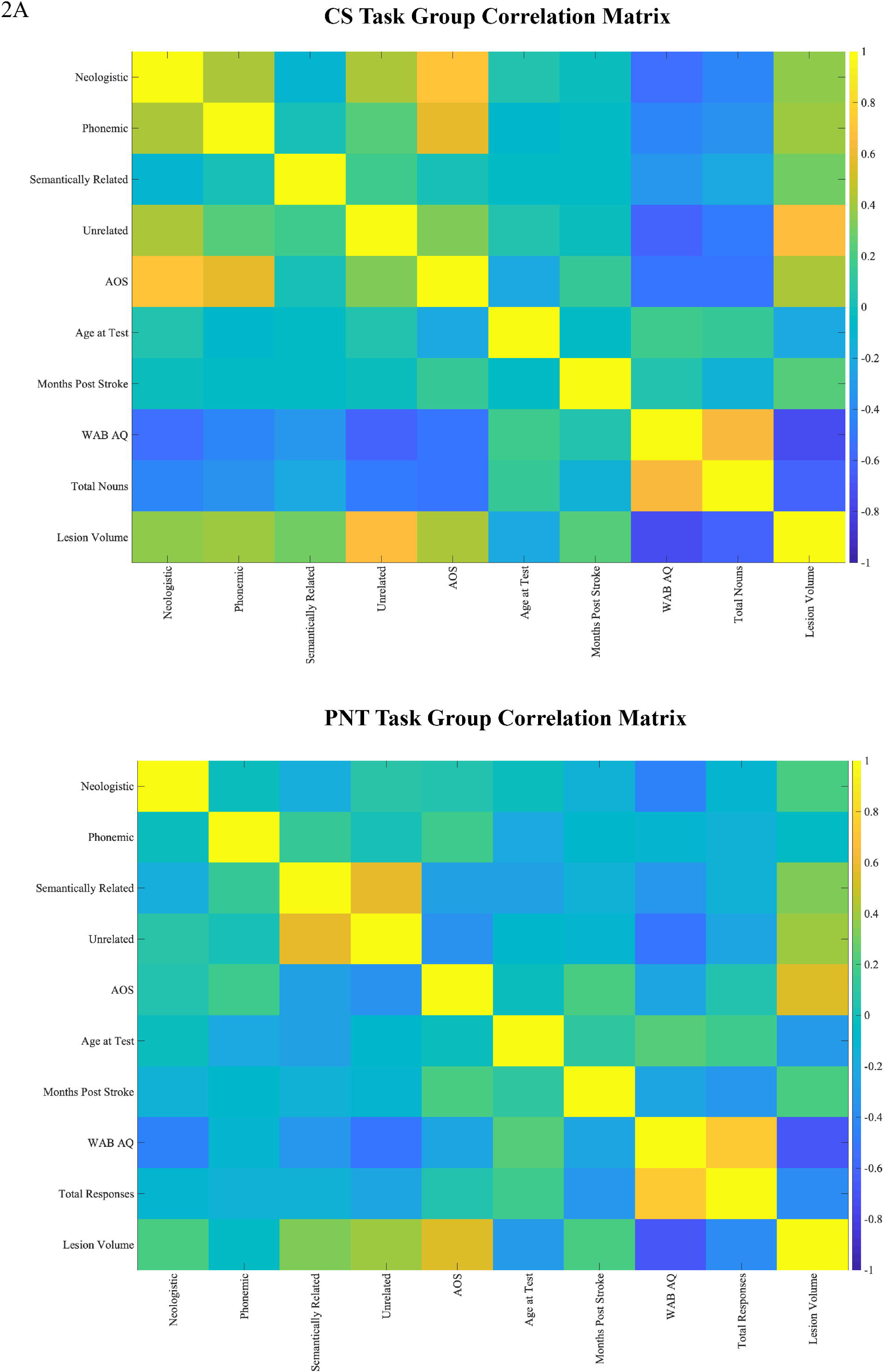
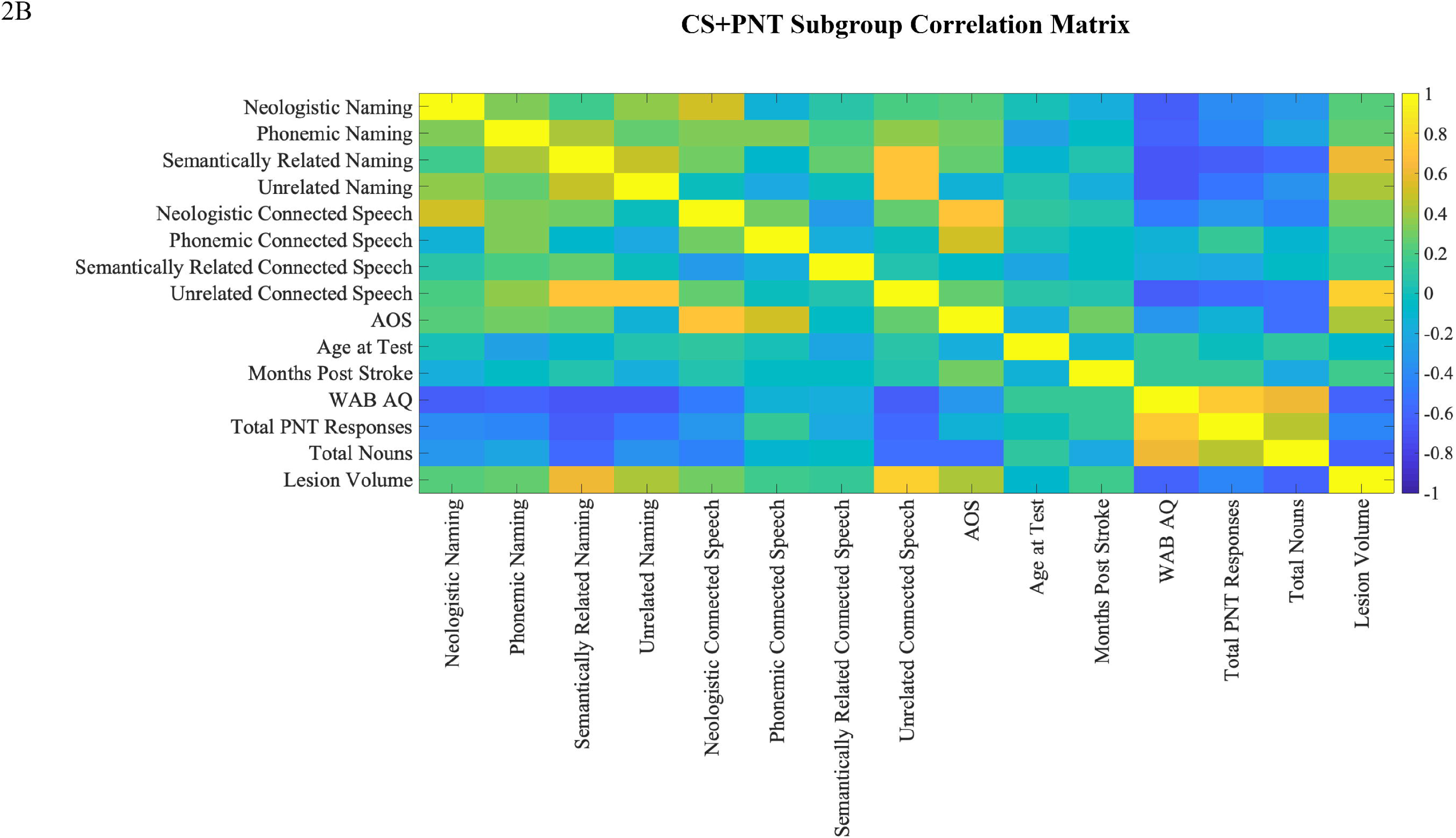
<u>Correlation matrices of groups</u>. A) CS Task Group, B) PNT Task Group and C) CS+PNT subgroup. Correlation matrices demonstrating relationships of biographical and linguistic variables with paraphasias by group. Cooler colors are strongly negatively correlated and warmer colors are strongly positively correlated. Matrices included paraphasias made during the respective task; biographical variables (age, time since stroke, lesion volume); and language variables (WAB AQ [Western Aphasia Battery aphasia quotient], total words spoken / total items named, apraxia of speech (AOS) score from the Apraxia of Speech Rating Scale (ASRS, Strand et al., 2014). Note that the CS task group showed significant correlations of paraphasia proportions (neologistic, phonemic, unrelated) with apraxia of speech severity, and this was why apraxia of speech severity was used as a covariate in VLSM analyses for the group.

The CS task group showed significant correlations between paraphasia proportions and apraxia of speech severity (neologistic, r=.71, p<.001; phonemic, r=.57, p<.001; unrelated, r=.32, p=.01). There were no significant correlations between any paraphasia type with age or months post-stroke (all r < .04). Both lesion volume (neologistic, r=.37; phonemic, r=.39; semantically related, r=.29; unrelated, r=.66) and AQ (neologistic, r=-.55; phonemic, r=-.42; semantically related=-.33; unrelated, r=-.62) were significantly related to paraphasia proportion (all p<0.05 after multiple comparison correction). Lesion volume was significantly related to AQ (r=-.73, p<.001). For subsequent VLSM analyses, lesion volume was used as a covariate, and for phonemic, neologistic and unrelated paraphasias, apraxia of speech severity was also used as a covariate.

The PNT task group did not show significant correlations between paraphasia proportions and apraxia of speech severity (all r < |.36|, all p>0.05). There were no significant correlations between any paraphasia type and age or months post-stroke (all r<|.28|, all p>0.05). Lesion volume significantly correlated with verbal paraphasias (semantically related, r=.34; unrelated=.41; all p<0.05 after multiple comparison correction) but not sound paraphasias (neologistic, r=.21; phonemic, r=-.05). AQ showed a similar pattern of correlation (neologistic, r=-.44; semantically related, r=-.32; unrelated, r=-.53; all p<0.05 after multiple comparison correction) but did not correlate with phonemic paraphasias (r=-.11). Lesion volume was significantly related to AQ (r=-.67, p<.001). For subsequent VLSM analyses, lesion volume was used as a covariate.

#### Comparison of Paraphasia Proportion Between Connected Speech and Naming

In the CS+PNT subgroup (N=36), comprised of individuals with scores for both naming and connected speech tasks, we were able to compare directly paraphasia proportions across tasks using multinomial logistic regression. Paraphasia proportions are shown in Table 2 and in the top portion of Figure 3.

**Figure 3:**
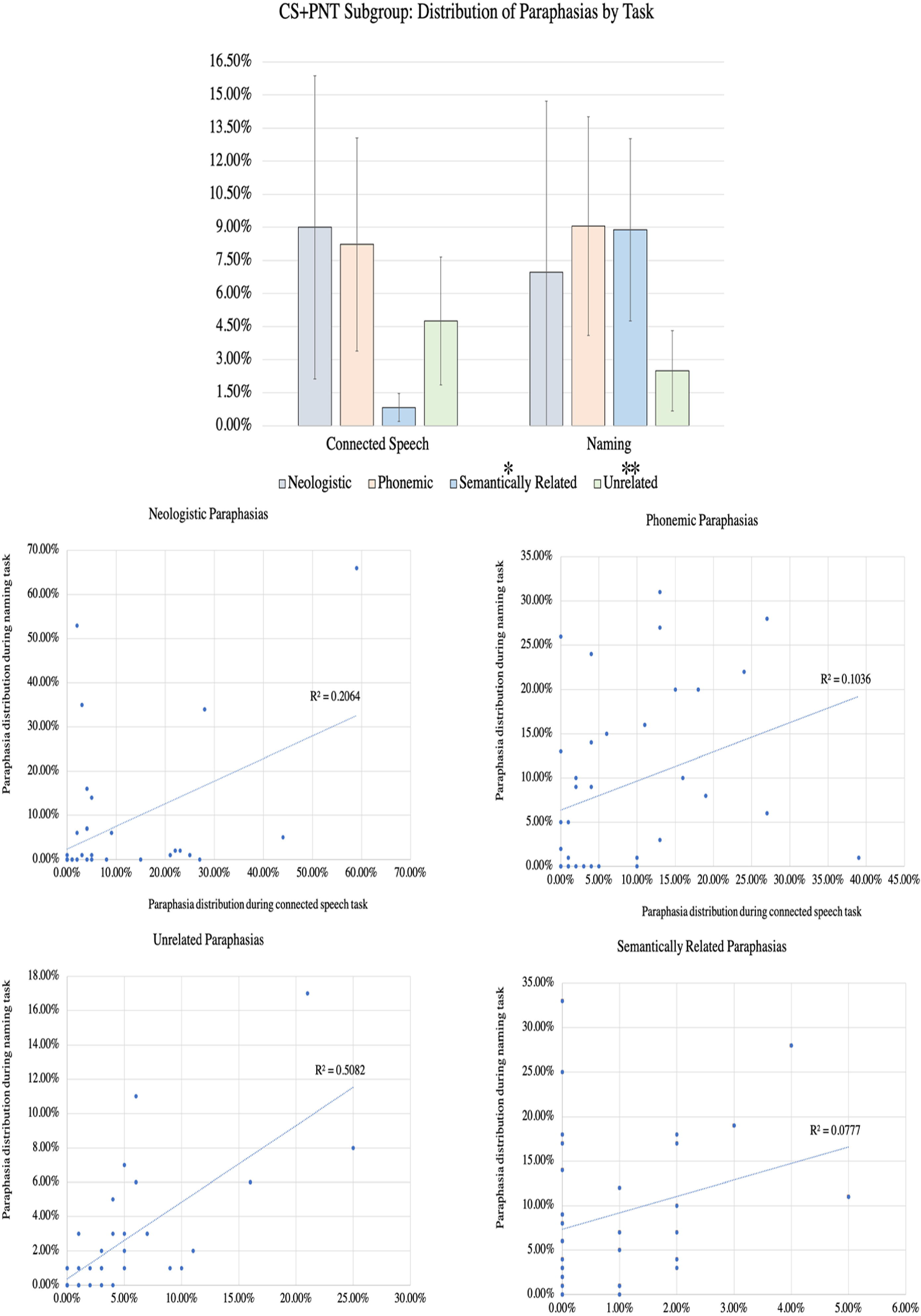
<u>Distribution of paraphasia types produced during connected speech and naming</u>. This figure directly compares the distribution of paraphasia types in the CS+PNT subgroup (N=36), the members of which completed both a connected speech and naming task. The top figure demonstrates the relative distribution of each paraphasia type by task. Multinomial logistic regression indicated that the semantically related and unrelated paraphasia types were significantly different between the tasks, and this is marked with a * (p<0.05) or ** (p<0.01) above the paraphasia name in the legend. The scatterplots (bottom) contrast the distribution during naming (y axis) and connected speech (x axis) for each paraphasia type.

A multinomial logistic regression was performed to model the relationship between paraphasia type (neologistic, phonemic, semantically related, unrelated) and task (connected speech, naming). Addition of the predictors to a model that contained only the intercept significantly improved the fit between model and data, χ^2^(4,N=72)=60.998, Nagelkerke R^2^=.76, p<.001. The parameter estimates showed a significant difference between tasks for two paraphasia types: semantically related (p=.001, Exp(B)=6.77^-49^) and unrelated (p=.015, Exp(B)=5.71^35^). Inspection of the data demonstrated greater proportion of semantically related paraphasias and fewer unrelated paraphasias during naming. The proportion of neologisms (p=0.13, Exp(B)=.001) and of phonemic paraphasias (p=0.48, Exp(B)=45.12) was not significantly different between tasks. Subsequent classification of paraphasia into type of assessment demonstrated an 88.9% correct classification into connected speech and 80.6% correct classification into naming task, with an overall accuracy of 84.7% (82.05% sensitivity [95% CI 66.47-92.46%] and 87.88% specificity [95% CI:71.80-96.60%]).

The scatterplots of the correlation between each paraphasia type by task are shown on the bottom of Figure 3.

### Voxelwise Lesion Symptom Mapping

#### Main Effect of Paraphasia Type

We evaluated brain damage significantly associated with each paraphasia type in the CS and the PNT task groups using VLSM. Figure 4A shows the VLSM results for each paraphasia type by task group, showing the voxelwise and cluster thresholded results. Table 3 serves as a complement to Figure 4A, presenting the maximum z-value and coordinates of voxels that, when damaged, significantly associated with paraphasia type by task group. Overall, for the CS task group, unrelated paraphasias were associated with a cluster of 14208 lesioned voxels in left hemisphere temporoparietal cortex; semantically related paraphasias were associated with a cluster of 19135 lesioned voxels in left hemisphere temporoparietal cortex; neologistic paraphasias were associated with a cluster of 14718 lesioned voxels in left hemisphere frontoparietal cortex and phonemically related paraphasias (when controlling also for apraxia of speech) were not associated with a significant cluster of damage. For the PNT task group, unrelated paraphasias were associated with a cluster of 19873 lesioned voxels in left hemisphere temporoparietal cortex and semantically related paraphasias were associated with a cluster of 20978 lesioned voxels in left hemisphere temporoparietal cortex.

**Table 3:**
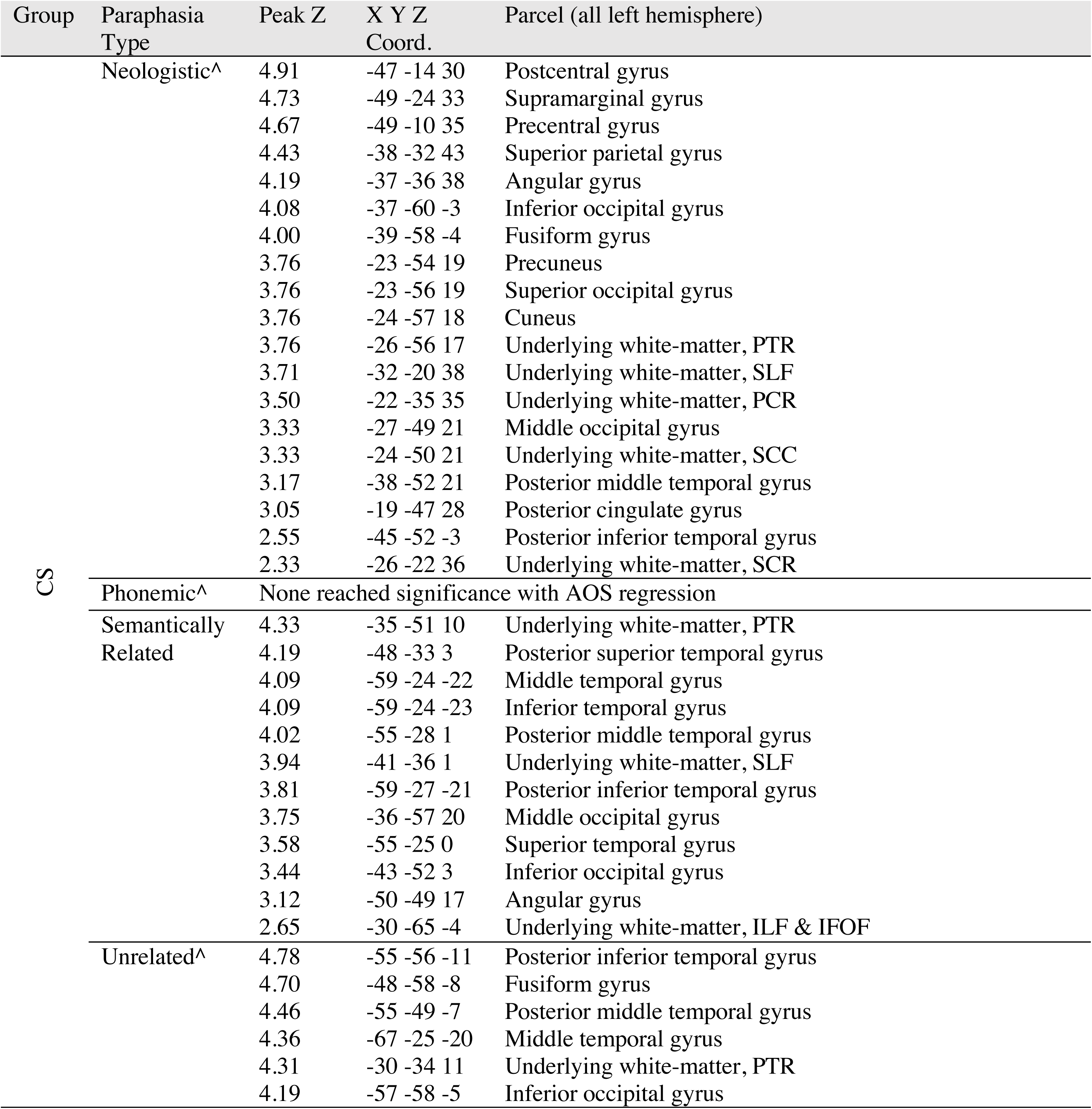
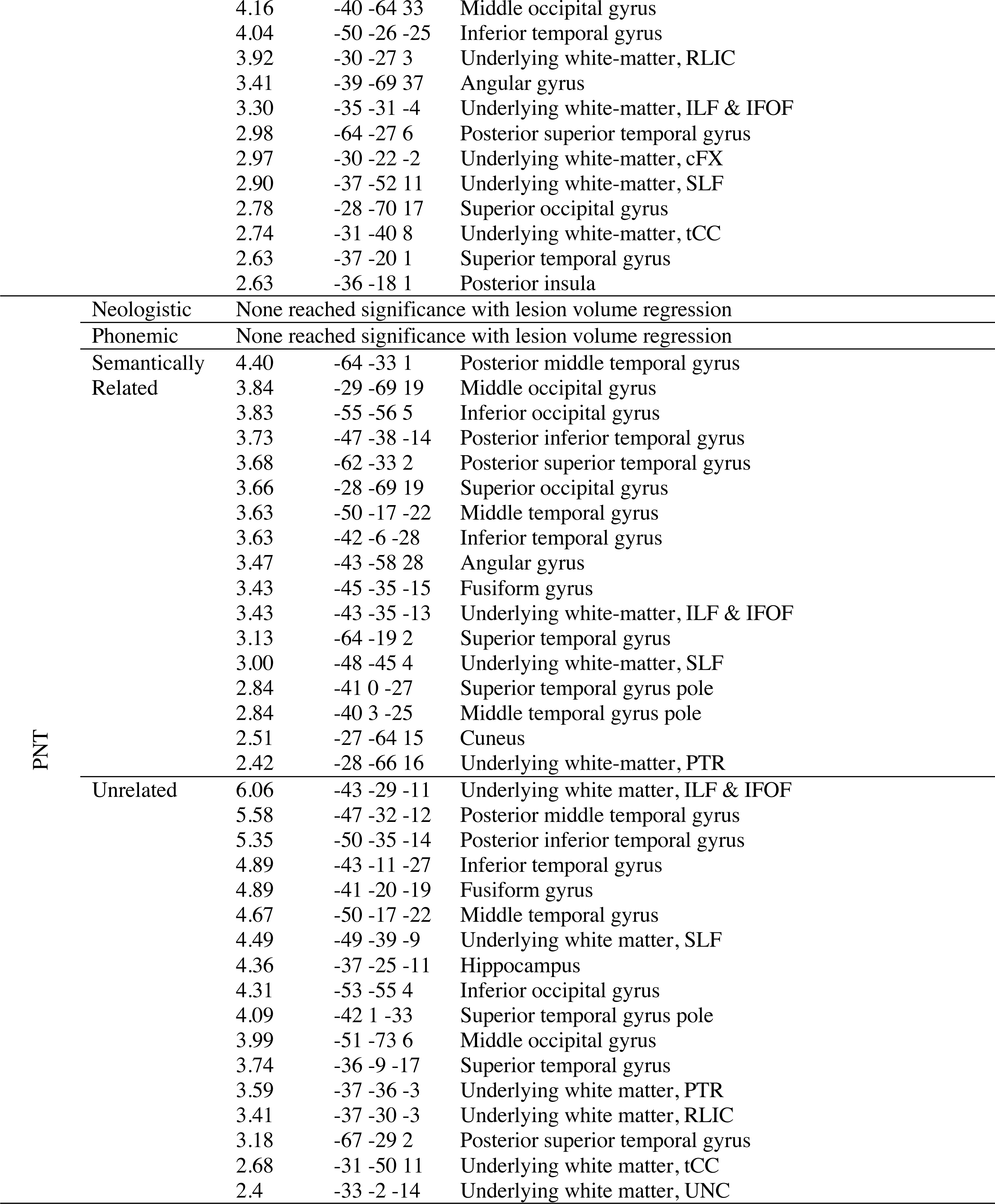
<u>Left hemisphere voxels that, when damaged, significantly correlate with paraphasia type by task group</u>. Here we present left hemisphere parcels containing voxels that, when damaged, significantly correlate with paraphasia type (neologistic, phonemic, unrelated, semantically related) for each task group (CS, PNT). To do this, we parcellated the lesion images using the Johns Hopkins University atlas, which includes gray and white matter (Faria et al., 2012). We then extracted the maximum z-score of voxels reaching significance within each parcel (“Peak Z” in table) and reported the coordinates of the maximum voxel (“X Y Z Coord.”). We report areas within the significant cluster, as described in the VLSM methods. All analyses included lesion volume as a covariate. This table serves as a complement to Figure 4A. ^ = analysis included both lesion volume and apraxia of speech (AOS) as covariate White matter abbreviations: cFX = fornix crescent; ILF = inferior longitudinal fasciculus; IFOF = inferior fronto-occipital fasciculus; PTR = posterior thalamic radiation; PCR = posterior corona radiata; RLIC = retrolenctiular part of internal capsule; SCC = splenium of the corpus callosum; SCR = superior corona radiata; SLF = superior longitudinal fasciculus; tCC = tapetum of the corpus callosum, UNC = uncinate Task group abbreviations: CS = Connected Speech Task Group (N=63); PNT = Naming Task Group (N=57)

**Figure 4:**
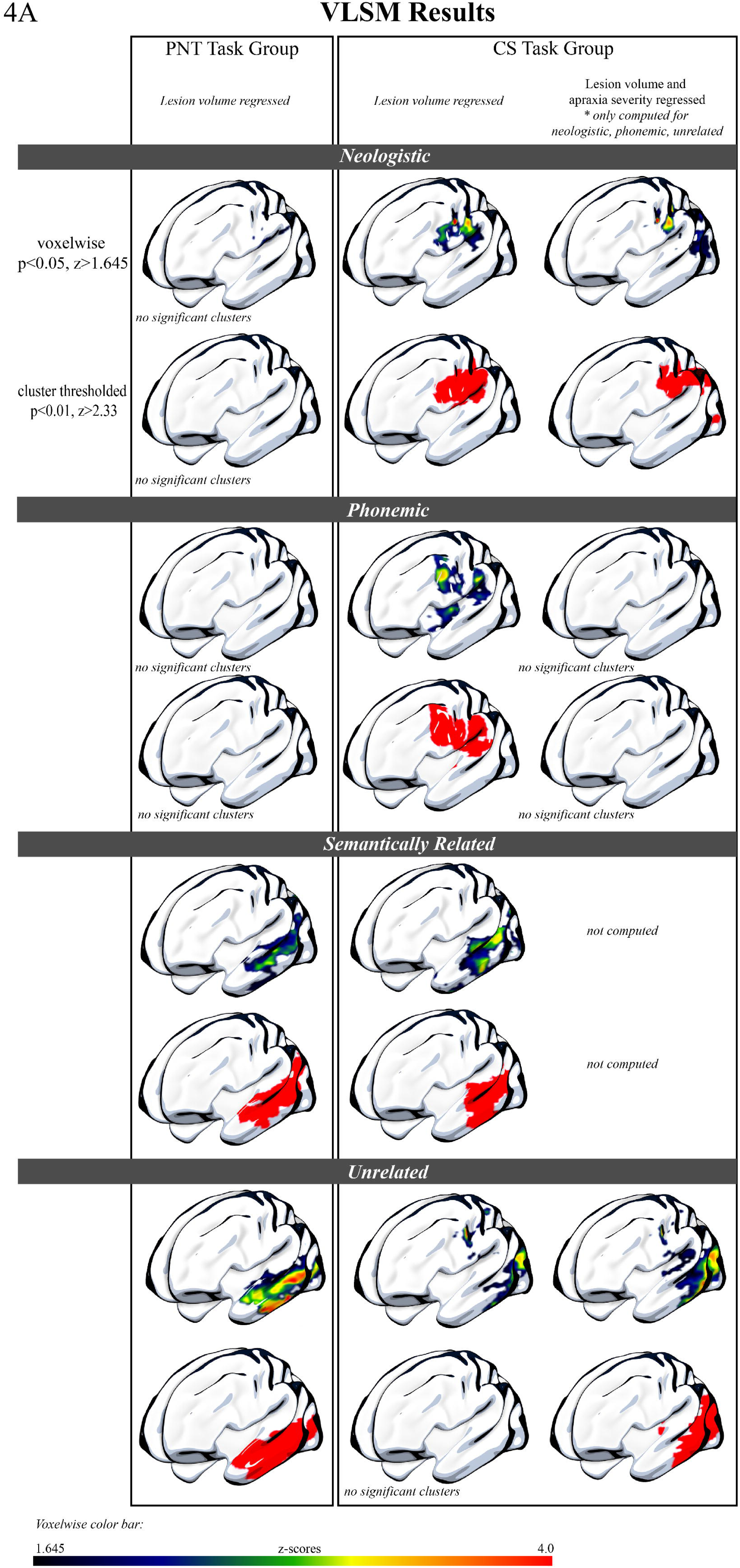
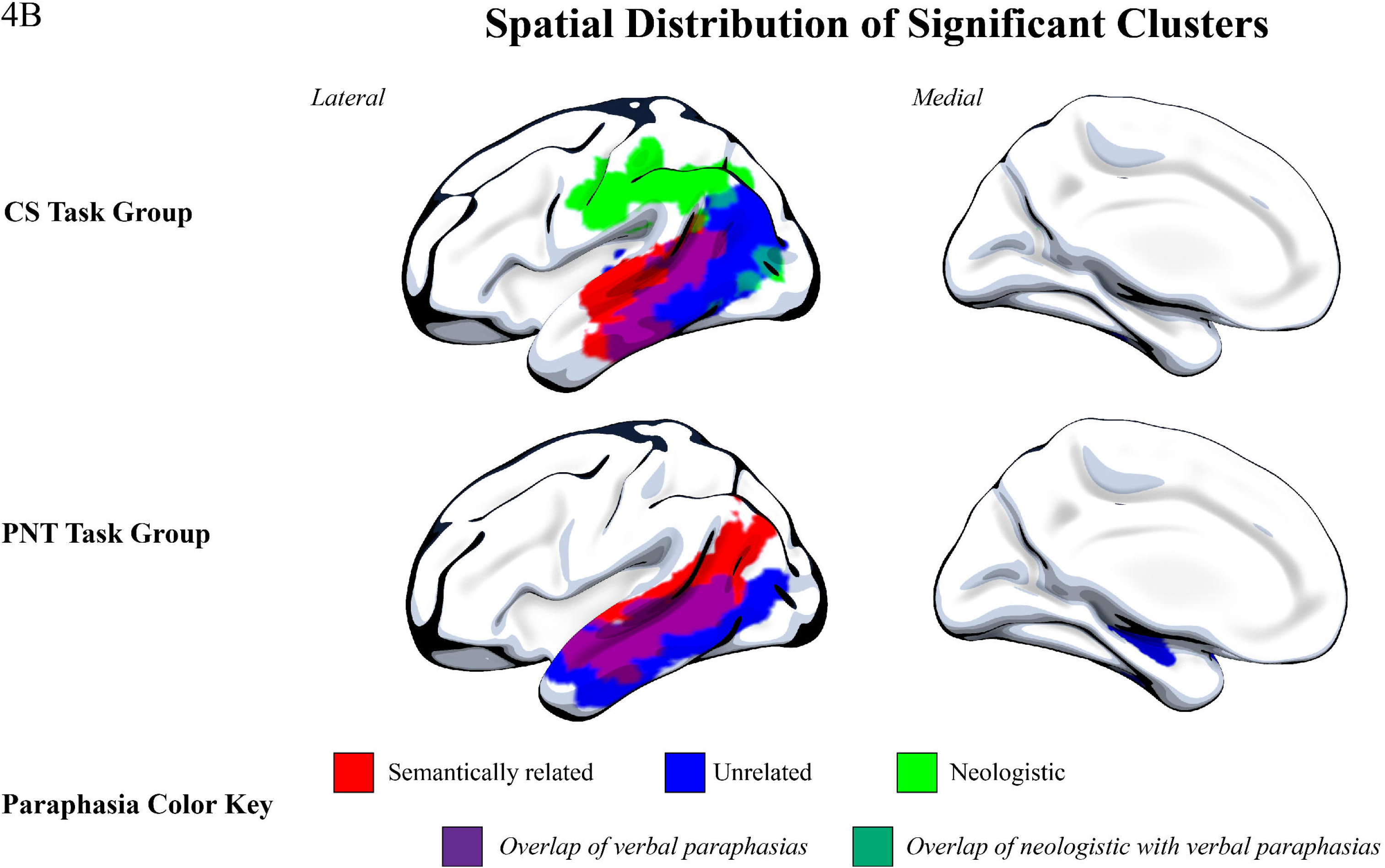
<u>Results of the voxelwise lesion symptom mapping on the CS and PNT task groups</u>. 4A: Surface rendering of the significant results of the VLSM analysis for the PNT task group (left) and CS task group (right). Both the PNT and CS task group results, which demonstrate damage significantly associated with each paraphasia type, have variance from lesion volume removed. In the far-right column, we also present results from the CS task group with variance from both lesion volume and apraxia of speech severity removed, as apraxia of speech severity was significantly correlated with neologistic, phonemic and unrelated paraphasias in this group. Lesion damage significantly associated with sound paraphasias (neologistic, phonemic) was only found for the CS task group, and demonstrated a dorsal distribution. Lesion damage significantly associated with verbal paraphasias (semantically related, unrelated) was found for both task groups, and demonstrated a more ventral distribution. 4B: Spatial distribution of significant clusters. Here, we demonstrate the spatial distribution of each significant cluster by paraphasia type and task group. As in Figure 4A, variance from lesion volume has been regressed from all analyses, and for the CS task group, we present neologistic and unrelated areas with apraxia of speech variance also regressed. Phonemic paraphasias are not demonstrated here, as there was not a significant cluster for phonemic paraphasia in the PNT task group after lesion volume regression or for the CS task group after both lesion volume and apraxia of speech severity regression. This figure demonstrates the distinction in dorsal and ventral streams associated with sound and verbal paraphasias, respectively.

Specifically, verbal paraphasias during connected speech were significantly associated with damage to left hemisphere ventral stream areas whereas sound paraphasias during connected speech were associated with damage to left hemisphere dorsal stream areas. Neologisms were significantly associated with damage to left hemisphere dorsal stream regions, including the supramarginal gyrus, postcentral gyrus, precentral gyrus, posterior superior temporal gyrus and underlying white matter. When both lesion volume and apraxia of speech severity were regressed, the pattern of damage associated with neologisms was similar. Phonemic paraphasias, when regressing variance from only lesion volume, were significantly associated with damage to more left hemisphere dorsosuperior areas as compared to neologisms, centering on supramarginal gyrus, precentral gyrus and middle frontal gyrus (posterior segment) but subsequent regression of apraxia of speech severity alongside lesion volume did not result in significant clusters for phonemic paraphasias. This is likely the result of sampling, where the sample with apraxia of speech also tended to present with more anterior lesion damage. Semantically related paraphasias were most associated with damage to left hemisphere posterior superior and middle temporal gyri, extending anteriorly into superior and middle temporal gyrus, and also included damage to angular gyrus and underlying white matter (superior longitudinal fasciculus, inferior longitudinal fasciculus, and inferior fronto-occipital fasciculus). Finally, no significant clusters were found to be associated with unrelated paraphasias when regressing only lesion volume; however, with the additional regression of variance from apraxia of speech severity, unrelated paraphasias were significantly associated with more posterior damage compared with semantically related paraphasias. These regions included damage to left hemisphere posterior superior, middle and inferior temporal gyri, angular gyrus and underlying white matter (superior longitudinal fasciculus, inferior longitudinal fasciculus, and inferior fronto-occipital fasciculus).

Results of the VLSM analysis for the PNT task group, looking at paraphasias types during naming, did not result in significant clusters of damage associated with sound errors (neologistic, phonemic). However, lesion damage associated with verbal paraphasias (semantically related, unrelated) was comparable to lesion damage associated with verbal paraphasias in the CS task group. Semantically related paraphasias significantly associated with damage to left hemisphere posterior superior, middle and inferior temporal gyri, extending anteriorly into middle, superior and temporal gyri and underlying white matter (superior longitudinal fasciculus). Unrelated paraphasias significantly associated with damage to left hemisphere posterior middle and inferior temporal gyrus and extending anteriorly into middle and inferior temporal gyri and underlying white matter (superior longitudinal fasciculus, inferior longitudinal fasciculus, and inferior fronto-occipital fasciculus).

As verbal paraphasias (semantically related, unrelated) were associated with a significant cluster of left hemisphere brain damage in both the CS and PNT task groups, we performed a conjunction analysis to analyze the extent to which these verbal paraphasias were associated with the same region(s) of the brain (Figure 5). Semantically related paraphasias significantly associated with a cluster of left hemisphere damage more anterior than unrelated paraphasias in the ventral stream, suggesting common neural mechanisms for these paraphasia types in naming and connected speech.

**Figure 5.**
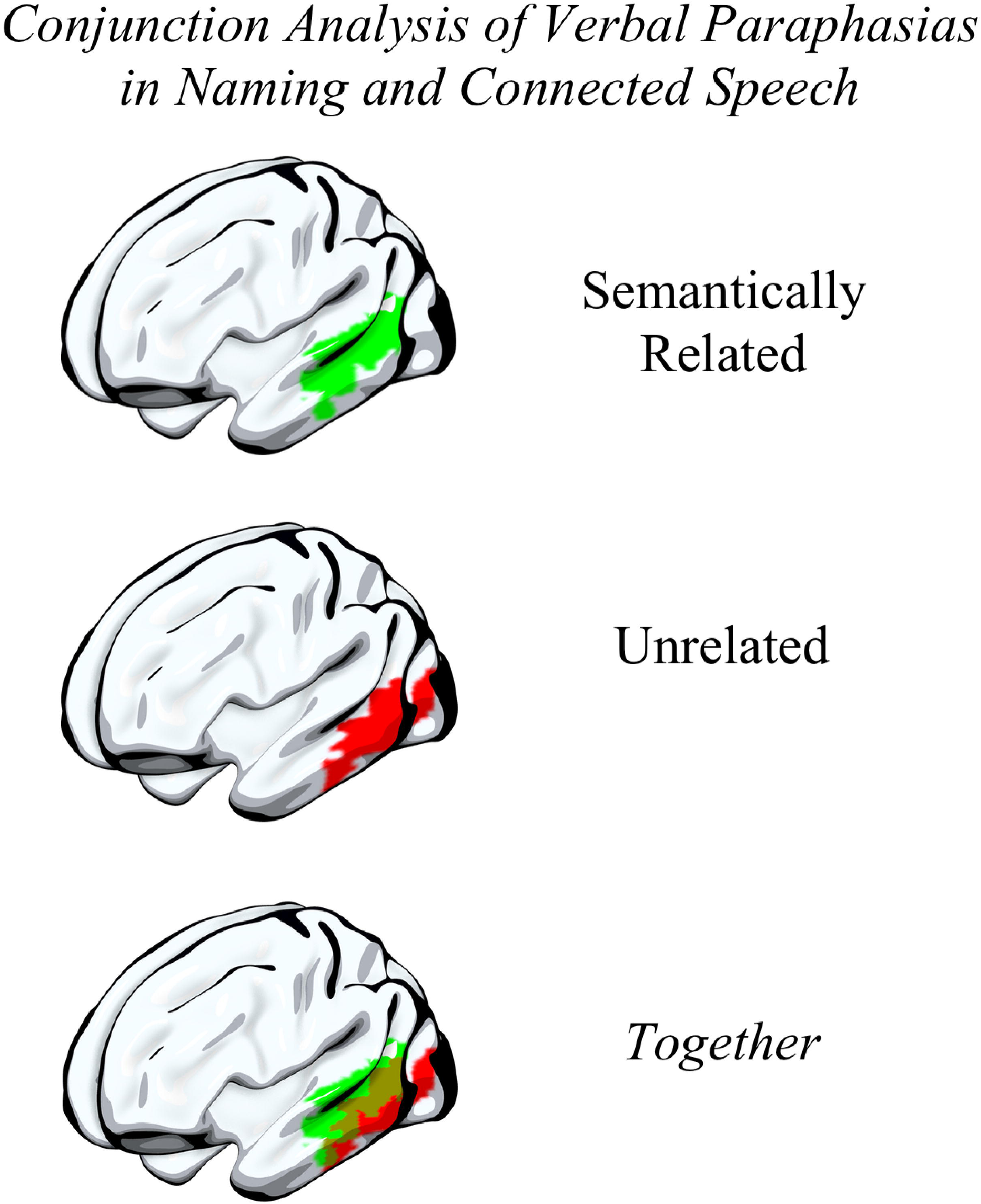
<u>Conjunction analysis of verbal paraphasias between tasks</u>. As verbal paraphasias in the connected speech and naming tasks significantly associated with lesion damage, we performed a conjunction analysis to identify the region(s) that were significantly associated with semantically related and with unrelated paraphasias in both tasks. A cluster of damage to more anterior, ventral stream areas, comprising superior and middle temporal gyri, was associated with semantically related paraphasias in both tasks (top) while a cluster of damage to more posterior, ventral stream areas was associated with unrelated paraphasias in both tasks (middle). There was an area of overlap in posterior superior and middle temporal gyri significantly associated with both semantically related and unrelated paraphasias in both tasks (bottom).

## Discussion

While prior studies associating brain damage with single-word errors have largely focused on paraphasias made during naming (single-word retrieval), very little work has focused on the neural substrates of connected speech in aphasia despite connected speech being a highly ecological and complex aspect of language that is strongly related to perceived quality of life (Cruice *et al.*, 2003; Mayer and Murray, 2003). Using a relatively large sample of post-stroke individuals, we were able to evaluate brain damage associated with paraphasias made during connected speech and naming and, in a sample of individuals who had scores on both a naming assessment and a connected speech assessment, compare the distribution of paraphasias across the tasks.

We first explored the differences in paraphasia proportion made during connected speech and naming. In the CS+PNT subgroup, we found that semantically related paraphasias occurred more often on the naming task than on the connected speech task, while unrelated paraphasias occurred more often on the connected speech task. An explanation for this finding is that the classification of verbal paraphasias in connected speech may be inherently difficult, as it is challenging to definitively know the intended target of a word when there are many possible targets. However, as demonstrated by the excellent inter-rater reliability on error coding across raters, this is unlikely to be the driving component behind the difference in the proportion of semantically related paraphasias across tasks. Further persuasive evidence toward the argument that verbal paraphasias were correctly classified in the connected speech task comes from the clear differentiation of lesion maps for each paraphasia type in both tasks (e.g. verbal paraphasias associated with ventral stream damage; sound paraphasias with dorsal stream damage) and the similarity in lesion maps for each paraphasia type across tasks (e.g. semantically related errors in both naming and connected speech associating with damage to left anterior temporal cortex). There are reasonable explanations for the difference in verbal paraphasia proportion across the tasks. Connected speech provides additional context in the form of prior word and future word selection, and the semantically related paraphasias that do occur during connected speech likely suggest a problem with semantic selection rather than a downstream problem of lexical selection. Therefore, the naming task may require higher lexical-semantic load than the connected speech task, a supposition that is supported by semantic interference demonstrated during blocked cyclic picture naming experiments (Schnur *et al.*, 2006; Oppenheim *et al.*, 2010). There is another possible explanation for the reduced proportion of semantically related paraphasias made during connected speech. During picture description, one can avoid talking about certain objects, whereas in picture naming, one is required to come up with a name for the picture. This may lead to a significantly smaller proportion of semantically related paraphasias in connected speech. Therefore, while there may be a reduced lexical-semantic load in connected speech, there are other strategies, such as circumlocution, that contribute to the reduced proportion of semantically related paraphasias made during connected speech. The overall greater proportion of unrelated paraphasias (always real words) and larger proportion of phonemic paraphasias and neologisms during connected speech also suggests an increase in lexical-phonological load, in keeping with the evidence proposing that several lexical items are selected simultaneously during connected speech (Kempen and Huijbers, 1983). An additional contributor to increased lexical-phonological load, aside from stress imposed by prior and future word choice and multiple word selection, is that connected speech is produced at a faster rate than the words produced during a picture naming task. Dell (1986) suggested that, when speech rate is increased, the available time for the spreading of activation across the language system is reduced, therefore activating incorrect items/sounds. In this way, previously activated items remain active because they have not had enough time to decay whilst new correct items have not received enough activation to compete at the same intensity. Indeed, we found that the number of neologisms made in the CS task group significantly correlated with the number of unrelated and phonemic paraphasias also made during connected speech whilst the number of neologisms made in the PNT task group did not significantly correlate with any other paraphasia type produced during naming. This postulates that lexical bias and phonemic similarity occur more often during connected speech (Oppenheim and Dell, 2008; Nozari and Dell, 2009; Oppenheim *et al.*, 2010).

Despite differing distributions of paraphasias across the two tasks, verbal paraphasias made during connected speech and naming were associated with damage to shared neural substrates (i.e. semantically related and unrelated paraphasias in connected speech and naming both associated with an area of damage localized to left hemisphere ventral stream cortex), supporting prior evidence from single-word retrieval that lesion damage in this area is associated with semantically related and unrelated paraphasias. While this result—that paraphasias in both connected speech and naming associate with areas of shared damage— seems straightforward and perhaps not novel, few studies have evaluated brain damage associated with paraphasias during connected speech (e.g. in primary progressive aphasia: Wilson *et al.*, 2010). As noted in the introduction, the two larger studies to do so focused on different populations (post-stroke aphasia; primary progressive aphasia) and also evaluated only one type of paraphasia (phonemic). In the present study, we find that verbal paraphasias, made during connected speech and naming, were associated with shared lesion damage to specific brain areas in the left hemisphere. Semantically related paraphasias have been long attributed to neurodegeneration (Jefferies and Lambon Ralph, 2006; Noppeney *et al.*, 2007) or stroke damage (Cloutman *et al.*, 2009; Schwartz *et al.*, 2009; Lambon Ralph *et al.*, 2012; Griffis *et al.*, 2017) involving left anterior temporal cortex, and this is reflected in our conjunction analysis which demonstrated that semantically related paraphasias made during both connected speech and naming localized to this area. Further, brain damage associated with unrelated paraphasias made during both tasks was localized to left posterior temporal cortex, an area which has been implicated most often in lexical retrieval (Lau *et al.*, 2008). These results further support the role of the left hemisphere ventral stream of language in lexical-semantic processing (Hickok and Poeppel, 2007).

Further, comparing lesion damage associated with verbal paraphasias made during connected speech allows us to make a finer distinction between semantic and unrelated errors in the anterior and posterior temporal lobe. Others have argued that semantic specificity increases on a gradient from posterior to anterior areas of the lateral temporal cortex, where the function of the anterior temporal lobe is most specific, especially specific for taxonomic and entity knowledge (for a review, see: Binder *et al.*, 2009). In people with post-stroke aphasia, semantic naming errors are most commonly taxonomic- or entity-specific (Jefferies and Lambon Ralph, 2006; Binder *et al.*, 2009; Walker *et al.*, 2011), where the resultant error is related by category, such as “apple” named as “pear". Semantically related paraphasias in our connected speech samples, which were likewise overwhelmingly taxonomic in nature, associated with more left anterior temporal lobe damage, suggesting subtle disruption of a more specific taxonomic- or entity-specific system. Whilst disruption in the left anterior temporal lobe results in a subtler impairment of lexical-semantic access (i.e. honing into the correct semantic form), damage to more posterior left temporal and occipital regions logically impairs lexical-semantic access by disrupting the lexical-semantic system at an earlier stage of retrieval (i.e. translating visual recognition). Supportive evidence for this comes from studies of focal damage to the left middle temporal gyrus, suggesting that this damage results in profound language comprehension and semantic deficits (Hart and Gordon, 1990; Hillis and Caramazza, 1991; Kertesz *et al.*, 1993; Dronkers *et al.*, 2004). We likewise showed that unrelated paraphasias made during connected speech—real words for which the target was unrelated or unknown—associated with left posterior temporal (especially middle and inferior temporal gyri) and left temporoparietal damage, indicating that disruption in areas recruited early on in the lexical-semantic system likely lead to verbal, but not phonological, paraphasias. Further, we demonstrate that damaged cortex in left posterior middle temporal gyrus area associated with semantically related and unrelated paraphasias made during both naming and connected speech, highlighting a role for this area of cortex in lexical selection. Our results, which evaluate verbal paraphasias for the first time in connected speech, support prior suppositions from single-word retrieval that the left posterior temporal lobe, in particular, has an early position in lexical-semantic access, at least during naming from visual recognition, and that the more anterior left temporal lobe, an area recruited downstream of the processes occurring in the left posterior temporal lobe, is an area whose role is more specific, likely for semantic entity knowledge.

Neologistic and phonemic paraphasias during connected speech associated with damage to left frontoparietal cortex. Neural models of motor speech control suggest that the left postcentral and supramarginal gyri are core areas for the mapping of somato-phoneme targets prior to production (Guenther and Vladusich, 2012; Hickok, 2012). As neologistic errors likely arise from phonological selection and/or substitutions, this result supports the supposition of the role of the dorsal stream in phonological retrieval (Hickok and Poeppel, 2007; Schwartz *et al.*, 2012). In general, few studies have evaluated brain damage associated with phonemic paraphasias in aphasia comparative to verbal paraphasias, and they have differed in their conclusions as to the loading of phonemic paraphasias onto the left hemisphere dorsal or the ventral stream of language (Kreisler *et al.*, 2000; Schwartz *et al.*, 2012). We found phonemic paraphasias during connected speech to be associated with frontoparietal left hemisphere brain damage. Notably, this result was only significant after removing variance from lesion volume but not apraxia of speech severity, suggesting a role for both word selection and motor planning in phonemic paraphasias produced during connected speech. Of particular interest is the robustness of VLSM results for sound paraphasias (neologistic, phonemic) derived during connected speech. In the present study, analyses performed on sound paraphasias from the connected speech task, and not the naming task, demonstrated significant results following removal of lesion volume variance and related apraxia of speech variance. Therefore, connected speech may be a particularly sensitive task on which to further evaluate lexical-phonological processing in the brain.

Our results extend prior behavioral research which described a divergence between paraphasias made during connected speech and naming (Nicholas *et al.*, 1989; Mayer and Murray, 2003; Fergadiotis and Wright, 2015) and our VLSM results establish a foundation for continued brain research comparing connected speech with naming. While we found that underlying brain structures are shared between paraphasias across tasks, at least as measured by lesion analysis, our behavioral analysis reflects subtle changes in linguistic load across tasks. While logically, a paraphasia should arise from damage to a brain area associated with the category of the paraphasia (i.e. left hemisphere anterior temporal lobe and semantically related errors), and therefore brain damage associated with paraphasias in connected speech and naming should be similar, there is a body of literature in neurotypical adults and adults with neurogenic communication disorders that connected speech involves the interaction of brain areas important for cognitive processes not usually found in single-word retrieval, such as inhibition and working memory (Coelho *et al.*, 1995; Wright *et al.*, 2014; Schnur, 2017). The multiword environment present during connected speech is prone to many processes not at play during single-word retrieval, including lexical bias (Nozari and Dell, 2009), phonemic similarity (Dell, 1984), and perhaps even semantic interference (Schnur *et al.*, 2006; Oppenheim *et al.*, 2010). We suggest and have first shown here that studying the neural substrates of paraphasias in connected speech share brain areas also associated with paraphasias of naming, but we further suggest that studying paraphasias in connected speech extends our understanding of the neurobiology of language. In the present study, we employed lesion-symptom methods largely focused on identifying the convergence and divergence of brain damage associated with paraphasias in both connected speech and naming, and we cannot fully elaborate on the recruitment of other linguistic or extra-linguistic areas to paraphasias in connected speech. Future work evaluating functional and structural connectivity may elaborate on the relative contribution of extra-linguistic cognitive processes to paraphasias produced during connected speech. In the current study, we analyzed a single naturalistic measure of connected speech, picture description, but we acknowledge that other forms of connected speech may be more naturalistic, such as story retelling and procedural descriptions, and encourage future work to evaluate these tasks in order to more comprehensively evaluate the spontaneous language system.

In conclusion, while analyzing connected speech comes with its own caveats, analysis of connected speech necessarily extends evidence garnered from single word retrieval and blocked cyclic picture naming to more naturalistic phenomena and necessarily extends our understanding of the language system and its neural underpinnings. Here, we demonstrate compelling evidence to pursue further behavioral and neuroimaging work in this area.

## Acknowledgments

We acknowledge the work of Kristie R Kannaley *MA MSP* in transcription and coding of speech samples.

## Funding

NIDCD P50 DC014664 [Fridriksson]

NIDCD U01 DC011739 [Fridriksson]

NIDCD R01 DC008355 [Fridriksson]

NIDCD T32 DC 014435 [Basilakos]

---

Supplementary Figure 1. <u>Flowchart of groups included in this study</u>. A retrospective sample of 120 participants was further classified into two mutually exclusive groups for use in the VLSM study: CS task group, which included participants with a connected speech assessment, and PNT task group, which included participants with a naming assessment. The PNT task group was matched, using case-control matching, on the aphasia severity of the CS task group. To directly evaluate behavioral distribution of paraphasias, we evaluated a subgroup of the CS task group, which we have called the CS+PNT subgroup because members of this group had both a connected speech and naming assessment. Notably, members of this group were not included in the PNT task group, as we created the CS and PNT task groups (for VLSM analysis) to be mutually exclusive.

