## Supplementary figures and images for "Neural organization of speech production: A lesion-based study of error patterns in connected speech"

### Supplementary Figure 1

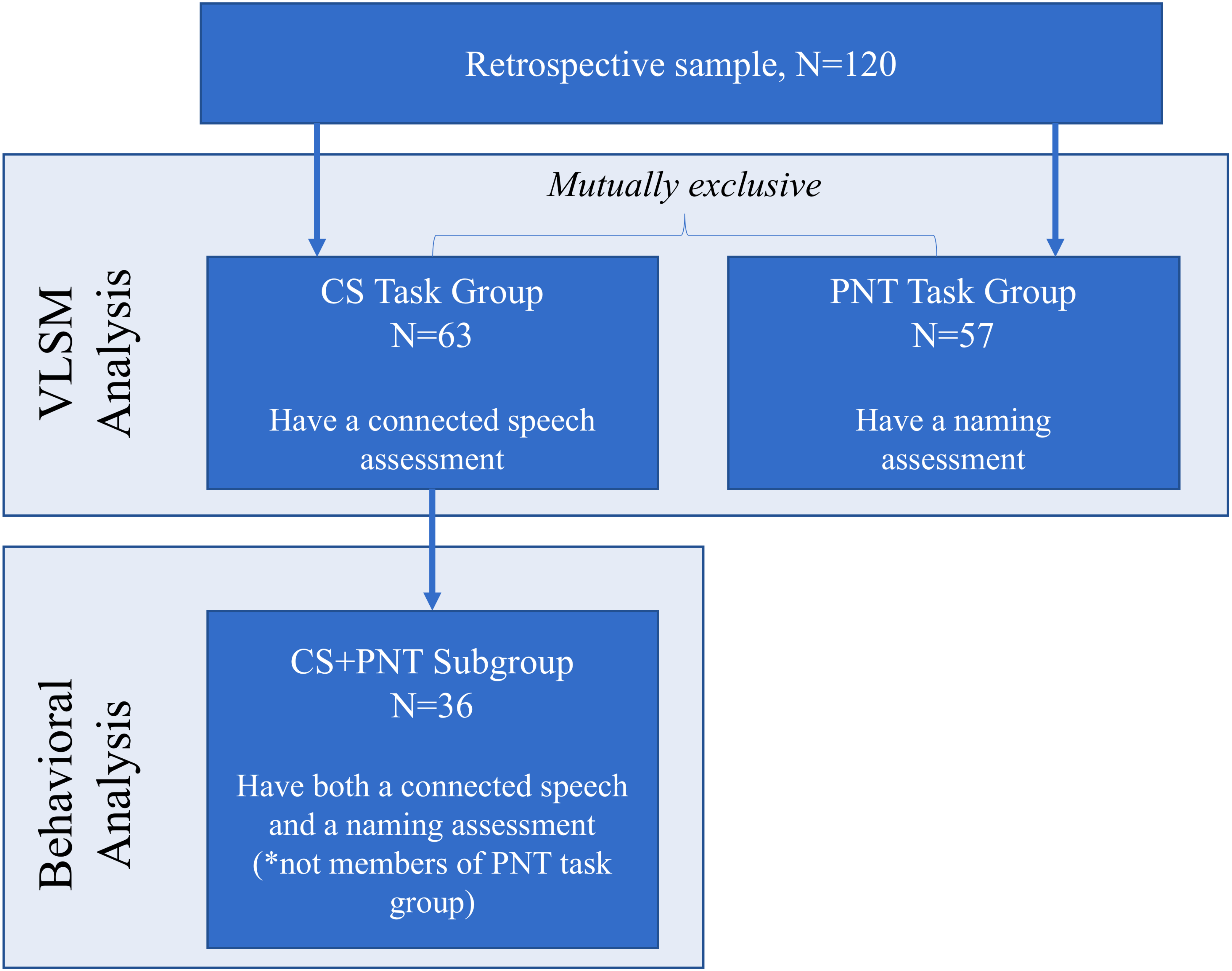
