## Supplementary Table 1 for "Neural organization of speech production: A lesion-based study of error patterns in connected speech"

Supplementary Table 1: Characterizing the targets of connected speech paraphasias. As the naming test was a test on nouns, the paraphasias made during the naming test replaced noun targets. To best compare the paraphasias made during connected speech and naming, we only used connected speech paraphasias in which the target replaced was a noun in the CSN group. The total number of paraphasias across all utterances, organized by type, is shown in the first row of data. The second row demonstrates the percentage of paraphasias which replaced noun targets (e.g. a paraphasia was made in place of a noun, rather than any other part of speech). Note that the target being replaced in neologisms and non-word phonemic paraphasias were more difficult to identify, as neologisms are non-words that do not share a substantial proportion of phonemes with a target.

| **Descriptives** | **All Paraphasias** | **Neologisms** | **Phonemic** | **Semantically Related** | **Unrelated** |
| --- | --- | --- | --- | --- | --- |
| **Total Number (M, SD)** | 472 (13.88±12.05) | 147 (4.32±6.01) | 214 (6.29±7.12) | 24 (0.71±0.97) | 87 (2.56±2.68) |
| **% of Noun Targets (SD)** | 57.35% (25.99) | 22.76% (27.97) | 57.62% (36.56) | 96.67% (12.91) | 74.46% (35.09) |
