## Supplementary Table 2 for "Neural organization of speech production: A lesion-based study of error patterns in connected speech"

Supplementary Table 2: Lesion proportion damage and the number of participants with damage to each left hemisphere gray- and white-matter parcel, organized by brain area. The table below includes every left hemisphere parcel (gray- and white-matter, Johns Hopkins University atlas) where at least 10% of subjects had damage. Divided by task group (PNT, CS), we then detail the proportion of damage (“Prop. Damage”) and the number of participants with damage (“N with Damage”) to each parcel.

|  | **Task Group** | **PNT** | | **CS** | |
| --- | --- | --- | --- | --- | --- |
|  | **Left hemisphere area** | **Prop. Damage** | **N with Damage** | **Prop. Damage** | **N with Damage** |
| Supra-Sylvian Areas | Superior frontal gyrus | 5.810168 | 30 | 3.372877 | 22 |
|  | Superior frontal gyrus prefrontal cortex | 1.350683 | 14 | 1.090516 | 11 |
|  | Middle frontal gyrus, posterior segment | 13.635578 | 39 | 7.068 | 31 |
|  | Middle frontal gyrus, dorsal prefrontal cortex | 4.139628 | 23 | 2.380625 | 21 |
|  | Inferior frontal gyrus, pars opercularis | 23.821532 | 42 | 17.928226 | 35 |
|  | Inferior frontal gyrus, pars orbitalis | 10.700935 | 35 | 10.774219 | 28 |
|  | Inferior frontal gyrus, pars triangularis | 18.248069 | 41 | 14.18063 | 30 |
|  | Lateral fronto-orbital gyrus | 5.129446 | 23 | 4.707771 | 27 |
|  | Middle fronto-orbital gyrus | 1.407397 | 6 | 0.401414 | 8 |
|  | Postcentral gyrus | 13.019269 | 50 | 8.623364 | 48 |
|  | Precentral gyrus | 16.325613 | 46 | 10.704902 | 44 |
|  | Superior parietal gyrus | 9.45292 | 43 | 8.965116 | 41 |
|  | Supramarginal gyrus | 21.417205 | 53 | 16.922147 | 46 |
|  | Angular gyrus | 17.855153 | 44 | 16.956421 | 41 |
|  | Precuneus | 1.272237 | 23 | 1.834755 | 20 |
|  | Anterior insula | 23.454689 | 45 | 20.611663 | 36 |
|  | Posterior insula | 29.141155 | 51 | 26.029184 | 43 |
|  | *MEAN (SD)* | *13.34 (8.26)* | *36.28 (13.35)* | *11.03 (7.61)* | *31.89 (11.99)* |
| Sub-Sylvian Areas | Superior temporal gyrus | 18.748946 | 46 | 24.1406 | 42 |
|  | Pole of superior temporal gyrus | 11.660901 | 40 | 15.962147 | 33 |
|  | Posterior superior temporal gyrus | 23.963974 | 45 | 25.989586 | 42 |
|  | Middle temporal gyrus | 8.675025 | 28 | 14.858871 | 33 |
|  | Pole of middle temporal gyrus | 4.933289 | 14 | 8.177952 | 19 |
|  | Posterior middle temporal gyrus | 14.956219 | 40 | 20.991522 | 40 |
|  | Inferior temporal gyrus | 4.124978 | 19 | 6.86684 | 27 |
|  | Posterior inferior temporal gyrus | 3.122858 | 19 | 7.228355 | 28 |
|  | Parahippocampal gyrus | 0.989051 | 9 | 1.177007 | 10 |
|  | Fusiform gyrus | 1.173892 | 17 | 1.694363 | 26 |
|  | Superior occipital gyrus | 4.568212 | 31 | 4.960698 | 32 |
|  | Middle occipital gyrus | 8.004182 | 36 | 9.837552 | 40 |
|  | Inferior occipital gyrus | 4.321997 | 21 | 6.95459 | 25 |
|  | Cuneus | 1.26895 | 20 | 0.697214 | 18 |
|  | Lingual gyrus | 0.987248 | 10 | 0.282534 | 11 |
|  | *MEAN (SD)* | *7.43 (7.03)* | *26.22 (12.56)* | *9.99 (8.53)* | *28.40 (10.53)* |
| White Matter | Corticospinal tract | 0.009104 | 13 | 0.009804 | 6 |
|  | Anterior corona radiata | 9.592864 | 39 | 7.684636 | 28 |
|  | Superior corona radiata | 24.034813 | 49 | 16.517966 | 47 |
|  | Posterior corona radiata | 21.216454 | 47 | 17.071486 | 46 |
|  | Genu of corpus callosum | 0.821429 | 23 | 1.116687 | 14 |
|  | Body of corpus callosum | 3.174786 | 30 | 2.526068 | 21 |
|  | Splenium of corpus callosum | 1.643651 | 30 | 1.604664 | 25 |
|  | Tapatum | 5.152276 | 23 | 5.131868 | 22 |
|  | Anterior limb of internal capsule | 8.952434 | 38 | 4.906367 | 30 |
|  | Posterior limb of internal capsule | 11.639974 | 40 | 9.257445 | 37 |
|  | Retrolenticular part of internal capsule | 16.136808 | 42 | 14.991159 | 43 |
|  | External capsule | 25.628165 | 51 | 19.41938 | 46 |
|  | Cingulum | 1.275675 | 23 | 1.695399 | 11 |
|  | Fornix / stria terminalis | 4.516543 | 25 | 0.35762 | 6 |
|  | Inferior fronto-occipital fasciculus | 14.512805 | 37 | 3.962188 | 27 |
|  | Posterior thalamic radiation | 9.752315 | 41 | 13.099464 | 32 |
|  | Sagittal stratum (including inferior longitudinal fasciculus and inferior fronto-occipital fasciculus) | 8.043796 | 38 | 8.88459 | 42 |
|  | Superior fronto-occipital fasciculus | 17.450122 | 34 | 9.69191 | 34 |
|  | Superior longitudinal fasciculus | 27.258592 | 56 | 12.615572 | 27 |
|  | Uncinate fasciculus | 11.198795 | 24 | 23.690686 | 55 |
|  | Ansa lenticularis | 6.829431 | 17 | 13.701807 | 26 |
|  | Lenticular fasciculus | 1.42492 | 6 | 3.879599 | 13 |
|  | *MEAN (SD)* | *10.47 (8.44)* | *33.00 (12.82)* | *8.72 (6.85)* | *29.00 (13.89)* |
| Medial Structures, including Basal Ganglia & Limbic System Structures | Amygdala | 3.491746 | 13 | 2.561781 | 13 |
|  | Hippocampus | 1.356514 | 10 | 1.840287 | 10 |
|  | Dorsal anterior cingulate gyrus | 2.091128 | 28 | 1.410309 | 19 |
|  | Posterior cingulate gyrus | 1.088063 | 27 | 1.078565 | 20 |
|  | Caudate | 2.358559 | 20 | 2.010155 | 19 |
|  | Putamen | 15.140834 | 48 | 10.20776 | 38 |
|  | Globus Pallidum | 8.522378 | 27 | 4.913986 | 22 |
|  | Thalamus | 1.643991 | 21 | 1.800241 | 25 |
|  | Mynert Nucelus | 2.133333 | 6 | 0.545455 | 7 |
|  | Cerebral Peduncle | 1.089827 | 5 | 0.337121 | 6 |
|  | *MEAN (SD)* | *3.89 (4.68)* | *20.50 )12.59)* | *2.67 (2.995)* | *17.90 (9.14)* |
